# Predicting Drug Protein Interaction using Quasi-Visual Question Answering System

**DOI:** 10.1101/588178

**Authors:** Shuangjia Zheng, Yongjian Li, Sheng Chen, Jun Xu, Yuedong Yang

**Author notes:** Co-first author (Shuangjia Zheng, Yongjian Li). Corresponding author (Jun Xu, Yuedong Yang).

## Abstract

Identifying novel drug-protein interactions is crucial for drug discovery. For this purpose, many machine learning-based methods have been developed based on drug descriptors and one-dimensional (1D) protein sequences. However, protein sequence can’t accurately reflect the interactions in 3D space. On the other hand, a direct input of 3D structure is of low efficiency due to the sparse 3D matrix, and is also prevented by limited number of co-crystal structures available for training. In this work, we propose an end-to-end deep learning framework to predict the interactions by representing proteins with 2D distance map from monomer structures (Image), and drugs with molecular linear notation (String), following the Visual Question Answering mode. For an efficient training of the system, we introduced a dynamic attentive convolutional neural network to learn fixed-size representations from the variable-length distance maps and a self-attentional sequential model to automatically extract semantic features from the linear notations. Extensive experiments demonstrate that our model obtains competitive performance against state-of-the-art baselines on the DUD-E, Human and BindingDB benchmark datasets. Further attention visualization provides biological interpretation to depict highlighted regions of both protein and drug molecules.

## 1 Introduction

Prediction of drugprotein interactions (DPIs) is of crucial importance for drug design and development. Though experimental assays remain to be the most reliable approach for determining DPIs, experimental characterization of every possible drug-protein pair is daunting due to the vast amount of money and labors in experiments.

Computational prediction of DPIs has therefore made rapid progress recently. In general, it falls roughly into two categories: physic-based and machine-learning method-s. Physic-based methods such as molecular docking apply physics-inspired predetermined energy functions to assess drug-protein interactions at the atomic level [Trott and Olson, 2010]. However, these methods are usually of limited accuracy due to difficulties to evaluate the conformational entropy and solvent contributions. Furthermore, these atom level methods are sensitive to structural fluctuations and can’t process protein flexibility well.

With the recent increase in protein structural data and protein-ligand interaction datasets, there is a rapid progress in machine learning-based methods [Ragoza *et al.*, 2017; Tsubaki *et al.*, 2018; Gao *et al.*, 2018]. Usually, the prediction is treated as a task of binary classification by integrating information of ligands, proteins, and their interactions in a unified framework.

Drug molecules can be well represented by their linear notations since most drugs contain less than 100 heavy atoms, and thus have a relatively small structural space. Recent studies have proven that current deep learning techniques can accurately predict structural properties from their linear representation [Zheng *et al.*, 2018; Öztürk *et al.*, 2018]. In contrast, protein molecules are much bigger, typically containing more than 1000 heavy atoms. The prediction from 1D sequence to 3D structure is the well-known challenging problem called protein folding. Therefore, traditional representation by 1D protein sequence is insufficient to capture the structural features in 3D space that decides the prediction of DPIs. Although the direct input of 3D structure was attempted in recent studies [Wallach *et al.*, 2015; Ragoza *et al.*, 2017; Stepniewska-Dziubinska *et al.*, 2018], they achieved relatively low accuracy due to a few reasons. First, the irregular protein 3D structure needs a big 3D matrix to contain the whole structure. The high-dimension, sparse matrix caused a large number of tedious input variables. Secondly, these studies suffered from small number of high-quality 3D structures because they need co-crystal structures of protein-ligand pairs that are difficult to determine by experiments.

As a balance, proteins can be alternatively represented by 2D pairwise distance map. Previous studies indicated that the distance map can recover protein 3D structures [Skolnick *et* al., 1997]. More recently, DeepMind group^1^ predicted protein structures more precisely than prior state-of-the-art so-lutions by using 2D image feature from a predicted distance map.

Inspired by these studies, we will utilize 2D distance map to represent proteins, and thus the DPI task can be converted into a classical Visual Question Answering (VQA) problem. Here, images are the distance maps for proteins, questions are the molecular linear notations for drugs, and answers are whether they will interact. This framework enables a training on protein monomer structures without need of co-crystal structures with their binding ligands, which significantly expands usable datasets for training.

However, there exists to be differences between VQA and DPI prediction. First, in many VQA scenarios, image size can be resized to a fixed value, but the distance map represents the real-world scale and can’t be resized. Second, the grammatical rules of chemical language are different from natural language, which force us to utilize customized tokenization process and suitable model for obtaining the semantic feature of molecular sequence. Third, our training set is still much smaller than other applications, which requires us to carefully design the networks.

To address the above problems, we present a VQA-inspired interpretable model that predicts DPIs directly from protein distance map and chemical language. The 2D map and chemical language are respectively encoded by a dynamic convolutional neural network (DynCNN) and bi-directional long short-term memory (BiLSTM) with attention, and the outputs are concatenated to dense layer to make prediction.

The proposed model is shown to outperform state-of-the-art approaches over three public DPI datasets. More importantly, the learned attentions enable visualized individual contributions between binding regions on proteins and ligands that is important for ligand refinement.

In summary, the main contributions of our work are as follows. To our best knowledge,

- this is the first study to utilize protein 2D distance map for predicting DPI;
- this is the first attempt to solve the DPI prediction with the VQA framework;
- extensive experiments are conducted on different level public datasets to demonstrate the effectiveness of our method.

## 2 Related works

Current drug-protein interaction prediction approaches could mainly be summarized as below:

**Docking-based** methods, such as [Trott and Olson, 2010; Koes *et al.*, 2013], are widely used to predict the binding mode and affinity given the 3D structure inputs of a drug compound and a protein. These methods apply predefined force fields to estimate the binding score to assess DPIs at the atomic level.

**Machine learning-based** methods have been investigated to predict DPIs. For example, [Bleakley and Yamanishi, 2009] proposed the bipartite local model by training local SVM classifiers from chemical structure similarity and protein sequence similarity. [Ballester and Mitchell, 2010] used random forest algorithm to capture binding effects during molecular docking process; [Durrant and McCammon, 2011] presented a scoring function based on fully connected neural networks to characterize the binding affinities of proteinligand complexes; [Tabei and Yamanishi, 2013] further improved the DPI prediction by using hashing algorithm with more compact fingerprints of compound-protein pairs.

Recently, deep learning techniques have been introduced to predict DPIs by direct use of 3D protein-compound complexes [Wallach *et al.*, 2015; Ragoza *et al.*, 2017; Stepniewska-Dziubinska *et al.*, 2018]. Since the input features were based on 3D matrix defined around pocket-ligand complexes, these methods generated a large number input variables, and suffered from limited number of training set.

In addition, though a few representation learning studies predicted DPIs based on protein sequence or their functional annotations [Tsubaki *et al.*, 2018; Gao *et al.*, 2018; Öztürk *et* al., 2018], their accuracies were limited with a lack of protein structural information.

## 3 Model

### 3.1 Problem Formulation

Our task is to predict the interaction between a drug compound and a target protein. Concretely, drug compound is represented in SMILES format, a text string for the topological information based on chemical bonding rules. For example, the benzene ring can be encoded as ‘c1ccccc1’. Each lowercase ‘c’ represents an aromatic carbon atom and ‘1’ for the start and closing of a cycle. All hydrogen atoms weren’t shown because they can be deduced via simple rules. To preserve important chemical features, we tokenized drug molecules using the following regular expression inspired by the work of [Olivecrona *et al.*, 2017]:

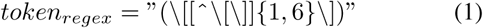

Additionally, we replaced the multi-character symbols using the following rules: ‘Br’:R, ‘Cl’:L, ‘Si’:A, ‘Se’:Z.

Suppose we have a drug molecular linear notation containing *n* tokens, the molecule can be represented in a sequence of molecular embeddings as *M* = (*t*_1_, *…, t*_*n*_), where *t*_*i*_ is a vector of *d*-dimensional token embedding for the *i*-th token. Thus, *M* ∈ ℝ^*n×d*^ is a representation of 2D matrix by concatenating all the token embeddings together.

Similarly, a protein can be simply described as a linear sequence that consists of a list of amino acids residues *P* = (*r*_1_, *…, r*_*l*_), where *r*_*i*_ is a one-hot representative vector with length of 20 for the amino acid type at position *i*, and *l* is the sequence length. Additionally, we calculate a 2D pairwise distance map by:

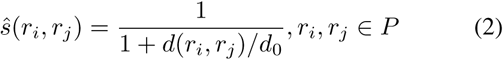

where *d*(*r*_*i*_, *r*_*j*_) is the distance between *C*_*α*_ atoms of residues *i* and *j*, and *d*_0_ is set to 3.8 *Å*. Let *ŝ*_*i*_ ∈ [*ŝ*(*r*_*i*_, *r*_1_), *…, ŝ*(*r*_*i*_, *r*_*l*_)] be the *l*-dimensional distance vector with all the residues in *P* of *r*_*i*_. The protein can be represented as a distance matrix:

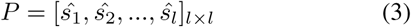

The goal of DPI prediction is to learn a system that takes a pair (*M*; *P*) as input and outputs label *y* ∈ {0, 1}, where *y* = 1 means an interaction between *M* and *P*.

Figure 1 is the architecture of our DrugVQA model. It comprises two main components: dynamic CNN with sequential attention (Sec.3.2) and BiLSTM with multi-head self-attention (Sec.3.3).

**Figure 1:**
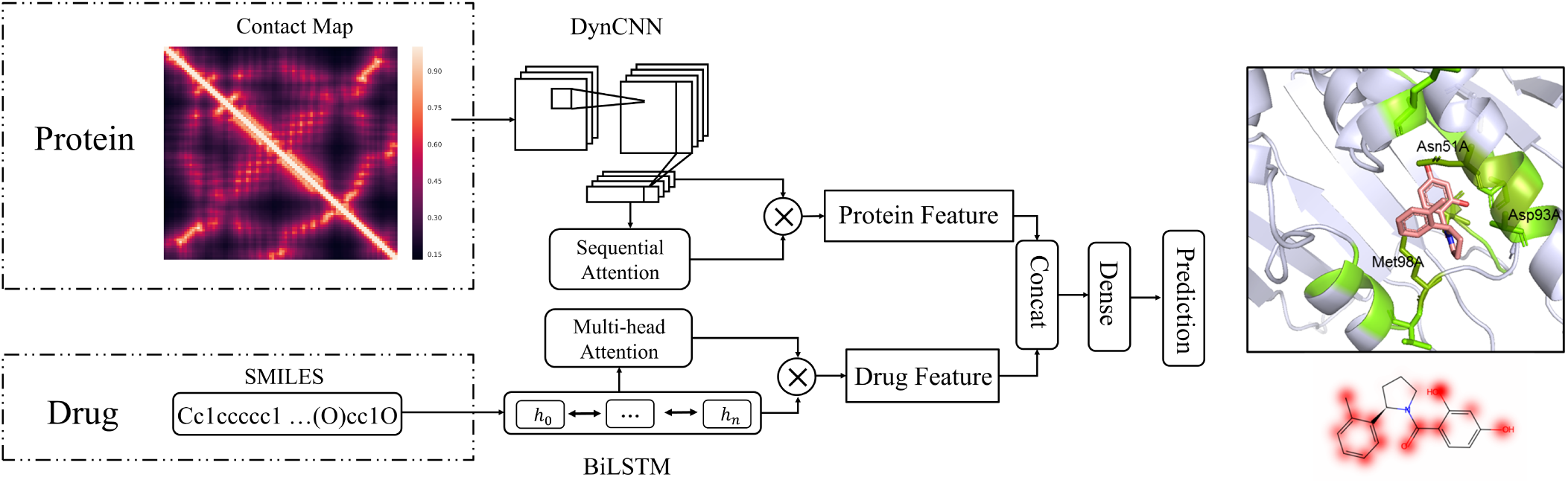
The framework of our proposed DrugVQA model. It consists of two main components, dynamic CNN with sequential attention and BiLSTM with multi-head self-attention.

### 3.2 Dynamic attentive CNN

#### Preliminary

In our model, adapted CNN is employed to code protein distance maps to fixed-size vector representations. Our CNN module consists of stacked residual blocks and a sequential self-attention block. For the residual block, we use modified Resnet [He *et al.*, 2016] to process protein inputs. Concretely, given a protein distance map *P* ∈ ℝ^*l×l*^, we apply a 3 × 3 convolutional layer with *N*_*f*1_ filters. The output is then processed with stacked residue blocks. The inputs of each intermediate residue units can be represented by *x*_*q*_, where *q*(1 ≤*q* ≤ *Q*) is the block unit index.

Each residue block could be defined as:

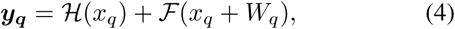

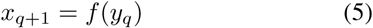

where *x*_*q*_ and *x*_*q*+1_ are input and output of the *q*-th block unit, and ℱ is a residual function, ℋ (*x*_*q*_) = *x*_*q*_ set as an iden-tity mapping. 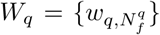 is a set of weights associated with the *q*-th residual unit, where 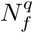 is the number of filters. The *f* function in Equation(5) is the activation function, and we utilize the Exponential Linear Unit (ELU) [Clevert *et al.*,2015] instead of traditionally used Rectified Linear Unit (ReLU). As shown in Figure 2(a), each residual unit is stacked by a 5 × 5 convolutional layer and a 3 × 3 convolutional layer.

**Figure 2:**
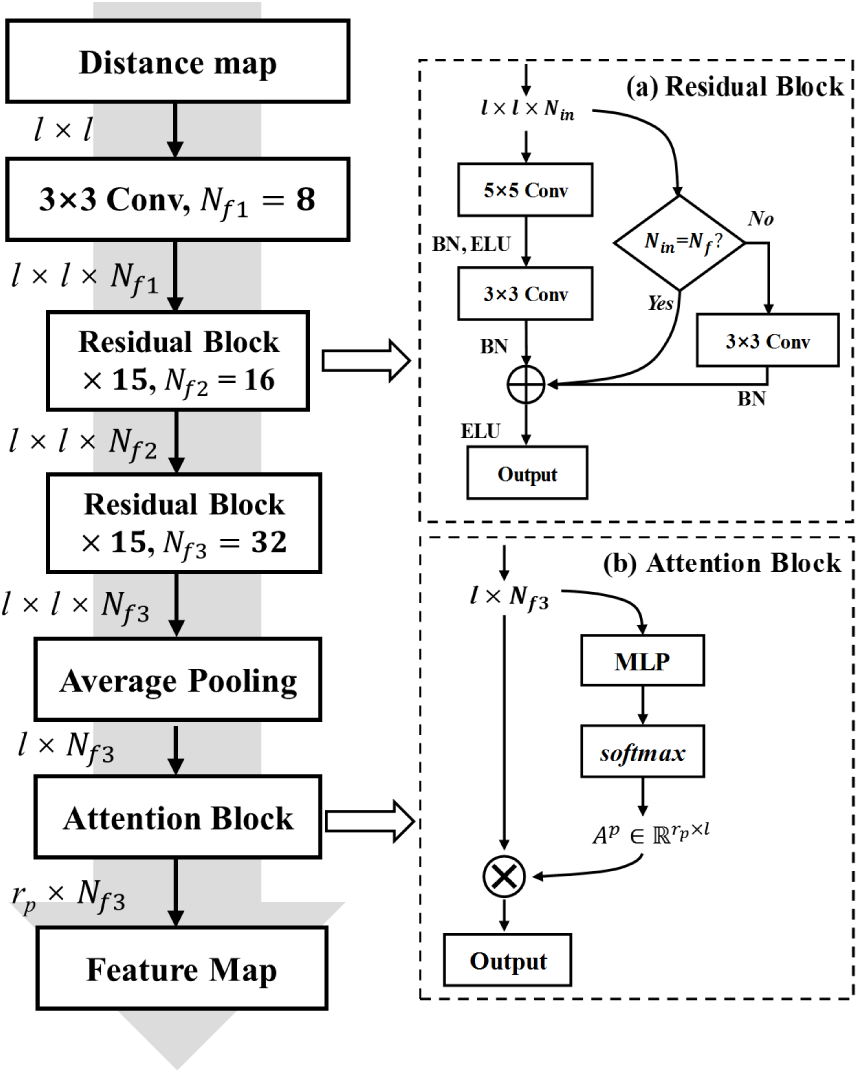
Dynamic attentive CNN. It includes two key components: (a) stacked residual blocks and (b) attention block.

#### Dynamic processing

Different from VQA tasks that often preprocess images to the same size, the real-world proteins are of different lengths of amino acids and can’t be scaled. Therefore, we want to design a dynamic neural network that could 1) handle inputs of variable lengths and 2) predict the importance of each amino acid. For this purpose, we take off the pooling layers between the residual block and use zero padding to two sides of input to ensure that the results of residual blocks have the same size as the input. Thus, the output of the last residual block remains the dimension of *l* × *l* × *N*_*f*_. Afterward, we apply average pooling to compress the information-enriched output of residual blocks for the downstream processing.

#### Sequential attention

Through average pooling, we obtain a protein feature map 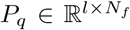. Practically, *P*_*q*_ can be viewed as protein sequential representation where *l* is the number of amino acids (sites) in the protein, and *N*_*f*_ represents the spatial feature of each site. As most sites are not directly related to the binding with drugs, recognizing the small portion binding sites is critical for the accurate prediction of DPIs. Inspired by the work of [Lin *et al.*, 2017], we adopt a multi-head sequential attention mechanism to fully use these features for classification. As shown in Figure 2(b), the attention mechanism takes the *P*_*q*_ as input, and outputs a vector of weights *a*^*p*^:

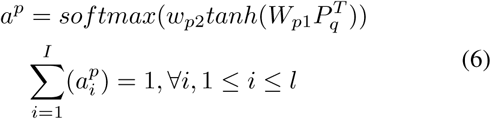

where 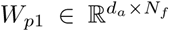, and *w*_*p*2_ is a vector of parameters with size *d*_*p*_, the *d*_*p*_ is an adjustable hyper-parameter. This vector representation usually focuses on a set of consecutive sites of protein sequence. Since a protein binding-pocket is composed of multiple consecutive sites neighbored in space, we further extend the *w*_*p*2_ into a *r*_*p*_-by-*d*_*p*_ matrix, noted as *W*_*p*2_, to capture the overall structural information of the binding-pocket. Thus, *a*^*p*^ is converted to a multi-head attention weight 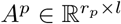as,

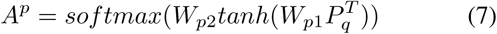

Practically, Equation (7) can be deemed as a 2-layer multilayer perceptron (MLP) without bias, whose hidden unit numbers is *d*_*a*_, and parameters are {*W*_*p*1_, *W*_*p*2_}. We compute the *r*_*p*_ weighted sums by multiplying the annotation matrix *A*^*p*^ and feature map *P*_*q*_:

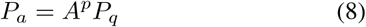

where *P*_*a*_ is an attentive feature map containing the latent relationship between contribution of sites on the interaction. The size of *P*_*a*_ is *r*_*p*_-by-*N*_*f*_.

### 3.3 Self-attentive BiLSTM

Each drug molecular SMILES string is encoded to a two-dimensional embedding matrix *M* ∈ ℝ^*n×d*^. Token vectors in the molecular matrix *M* are independent to each other. To gain some dependency between adjacent tokens within a molecule, a bi-directional LSTM is used to process a molecule:

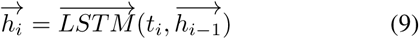

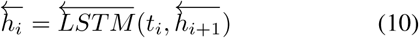

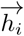 is concatenated with 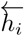, and a hidden state *h*_*i*_ is obtained to replace token embedding *t*_*i*_, and thus *h*_*t*_ becomes a more information-enriched vector that gains some dependency between adjacent tokens in a molecule. For simplicity, we note all *h*_*i*_ in every time step *i* as *H*.

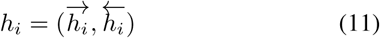

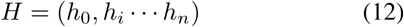

If the hidden unit number for each uni-directional LSTM is set as *u*, the shape of *H* would be *n*-by-2*u*.

The next goal is to know which part of the molecule contributed most to the interaction prediction. In other words, we want to identify the relationship between tokens and interaction, which can be used for a chemist to design or improve chemical compounds. Similarly, we achieve this by introducing multi-head self-attention mechanism. The attention mechanism takes the whole LSTM hidden states *H* as input, and outputs a vector of weights *A*^*m*^ as

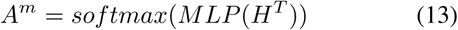

where hidden unit numbers of MLP is *d*_*m*_, and parameters are {*W*_*m*1_, *W*_*m*2_}. We compute the weighted sums by multiplying the annotation matrix *A*^*m*^ and LSTM hidden states *H*, the resulting matrix is the self-attentive molecular embedding:

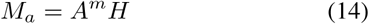

where *M*_*a*_ is a self-attentive drug molecular feature map that contains the latent relationship between tokens contribution of interaction. The size of *M*_*a*_ is *r*_*m*_-by-2*u*, where *r*_*m*_ is an adjustable hyper-parameter representing the number of attention vectors.

### 3.4 Classifier

For *P*_*a*_ and *M*_*a*_, we summed up over all the attention vectors, and then normalized the resulting weight vector to sum up to 1. This process enables us to obtain two information-enriched 1-D vectors 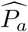 and 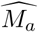, which will be fed into the classification layer. We concatenate 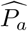 and 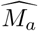, i.e., 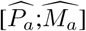, and obtain an output vector *o ∈* ℝ^2^, which is the input to DPI classifer:

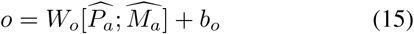

where 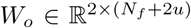 is the weight matrix and *b*_*o*_ ∈ ℝ^2^ is the bias vector. Finally, a *sigmoid* function is appended on top of the output layer *o* = [*y*_0_, *y*_1_] to model the DPI probability as follows:

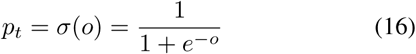

where *t* ∈ {0, 1} denotes the binary label (i.e., interact or not) and *p*_*t*_ is the probability of *t*, and we denote *ŷ* as the probability of *t* = 1.

### 3.5 Training

Given a dataset *D* = {(*m*_*i*_, *p*_*i*_, *y*_*i*_)}, the training objective is to minimize the loss function ℒ, given as the cross-entropy loss as follows:

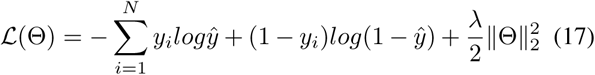

where Θ is the set of all weight matrices and bias vectors in our system, and *N* is the total number of drug-protein pairs in the training dataset, and *λ* is an L2 regularization hyperparameter. Θ is trained using the backpropagation algorithm.

## 4 Experiments

### 4.1 Dataset

To enable head-to-head comparisons of DrugVQA to existing machine learning-based methods and docking programs, we evaluated our proposed model on three public DPI datasets: the DUD-E dataset, the Human dataset, and BindingDB dataset.

#### DUD-E

The DUD-E is a well-known benchmark consisting of 102 targets across 8 protein families [Mysinger *et al.*, 2012]. On average, each target has 224 actives and over 10,000 decoys. Computational decoys are chosen such that they are physically similar but topologically dissimilar to the actives. The finally dataset contains 22, 645 positive examples and 1, 407, 145 negative examples. We adopt a three-fold cross-validation strategy to train and evaluate our model on the DUD-E dataset following [Ragoza *et al.*, 2017]. The folds were split between targets, where all ligands of the same target belong to the same fold. To avoid the impact of homologous proteins, targets belonging to the same protein families were strictly kept in the same fold. For a fast training of models, we used balanced set (all positives and randomly chosen equivalent negatives for each target) for training, but kept using the whole set (unbalanced ones) for evaluation.

#### Human

Created by [Liu *et al.*, 2015], this dataset includes highly credible negative samples of compound-protein pairs obtained by using a systematic screening framework. Following [Tsubaki *et al.*, 2018], we use a balanced dataset, where the ratio of positive and negative samples is 1:1. Finally, the human dataset contains 6, 675 interactions and 1, 998 unique proteins. We adopt the same five-fold cross validation strategy as in the original paper.

#### BindingDB

We further choose the BindingDB datasets [Gilson *et al.*, 2015] as the real-world dataset to evaluate our model. BindingDB is a public database of experimentally measured binding affinities, focusing chiefly on the interactions of small molecules and proteins. In our experiments, we use the customized BindingDB dataset constructed by [Gao *et* al., 2018] for head-to-head comparisons. The dataset contains 39, 747 positive examples and 31, 218 negative examples from bindingDB. We report the results on their customized testing datasets.

### 4.2 Implementation and Evaluation Strategy

#### Proposed Model

We implemented the proposed model with Pytorch 0.4.0 [Paszke *et al.*, 2017]. The training process lasts at most 50 epochs on all the datasets using the Adam optimizer with a learning rate of 0.001 and batch size of 1. Considering the limitation of memory of the used GPU (GTX1080Ti 12GB), we employed 30 residual blocks with 16 and 32 filters (the *N*_*f*2_ and *N*_*f*3_ in Figure 2), respectively. The hidden state of BiLSTM was set to 64 (the *u* in Sec 3.3), 0.2 dropout was applied on the BiLSTM and self-attention MLP unit. In addition, attention MLPs both in CNN and BiLSTM had a hidden layer with 100 units (the *d*_*p*_), and we chose the matrix embedding to have 10 rows (the *r*_*p*_) for protein and 18 rows (the *r*_*m*_) for drug. The coefficient of L2 regularization was 0.001. We explored hyperparameters in a wide range and find the above set of hyperparameters yields the highest performance.

#### Evaluation Metrics

Performance were evaluated by the area under the receiver operating characteristic curve (AUC). In addition, for Human dataset, we report the Precision and Recall value following [Tsubaki *et al.*, 2018]. For DUD-E dataset, we report the ROC enrichment metric (RE) following the work of [Ragoza *et al.*, 2017]. Specifically, the RE score is defined as the ratio of the true positive rate (TPR) to the false positive rate (FPR) at a given FPR threshold. Here, we report the RE scores at 0.5%, 1%, 2%, and 5% FPR thresholds. For BindingDB dataset, we also report the accuracy following [Gao *et al.*, 2018].

### 4.3 Comparisons on the Human dataset

#### Compared Models

In this section, we compare our DrugVQA with the state-of-the-art DPI approaches on the Human dataset. We compare it with k-NN, random forest (R-F), L2-logistic (L2), SVM models (results obtained from [Liu *et al.*, 2015]), and GNN [Tsubaki *et al.*, 2018] (we retrained the model with the same parameter settings as in the original papers). To verify the effectiveness of our proposed model, we also include a version of our model by replacing protein distance map with protein sequence (DrugVQA(seq)). The protein sequences were processed as drug SMILES through self-attentive BiLSTM.

#### Results

As shown in Table 1, DrugVQA outperforms protein sequence-based model with an increase of 1.9%, 3.3%, and 4.4% on AUC, recall, and precision, respectively. This agrees with our expectation that the distance maps contain more information than sequences for the prediction of DPI. The DrugVQA (seq) is comparable to GNN method though GNN employed graphical neural network for coding chemical structure of drugs. Due to relatively simple structure of drugs, the GNN doesn’t bring significant changes compared to our self-attentive BiLSTM. Other descriptor-based machine learning techniques have low performance with AUC ranging from 0.86 to 0.94, indicating that the end-to-end learned representations can learn important information from proteins and drugs for DPI prediction.

**Table 1:** Comparison results of proposed models and baselines on Human Dataset.

| Method | AUC | Recall | Precision |
| --- | --- | --- | --- |
| k-NN | 0.860 | 0.927 | 0.798 |
| RF | 0.940 | 0.897 | 0.861 |
| L2 | 0.911 | 0.913 | 0.861 |
| SVM | 0.910 | 0.950 | **0.966** |
| GNN | 0.970 | 0.918 | 0.923 |
| DrugVQA(seq) | 0.968 | 0.930 | 0.911 |
| DrugVQA | **0.987** | **0.963** | 0.955 |

### 4.4 Comparisons on the DUD-E dataset

#### Compared Models

We compare our DrugVQA with the state-of-the-art DPI approaches on DUD-E dataset, which can be divided into three categories: 1) conventional docking approaches Vina [Trott and Olson, 2010]; Smina [Koes *et* al., 2013]; 2) machine-learning scoring functions NN-Score [Durrant and McCammon, 2011]; RF-Score [Ballester and Mitchell, 2010]; 3) deep learning-based method 3D-CNN [Ragoza *et al.*, 2017]; AtomNet [Wallach *et al.*, 2015]; GNN [Tsubaki *et al.*, 2018].

#### Results

As listed in Tab 2, DrugVQA achieved an order-of-magnitude improvement over baselines at a level of accuracy useful for drug discovery. On the full DUD-E dataset, DrugVQA outperforms the state-of-the-art GNN model with an average AUC of 0.971 versus 0.94. Though 3D-CNN have employed three-dimensional structure for training, it has the lowest performance among three deep learning methods. This is likely due to the sparse data in 3D space, whereas the 2D pairwise distance map provides a good balance. As a result, DrugVQA outperforms 3D-CNN for 96% of the DUD-E targets on a per-target basis.

**Table 2:** Mean AUC and ROC Enrichment (RE) across targets on the DUD-E Dataset for proposed models and baselines.

| Model | AUC | 0.5% RE | 1.0% RE | 2.0% RE | 5.0% RE |
| --- | --- | --- | --- | --- | --- |
| Vina | 0.716 | 9.139 | 7.321 | 5.811 | 4.444 |
| Smina | 0.696 | - | - | - | - |
| NN-score | 0.584 | 4.166 | 2.980 | 2.460 | 1.891 |
| RF-score | 0.622 | 5.628 | 4.274 | 3.499 | 2.678 |
| 3D-CNN | 0.868 | 42.559 | 26.655 | 19.363 | 10.710 |
| AtomNet | 0.895 | - | - | - | - |
| GNN | 0.940 | - | - | - | - |
| DrugVQA | **0.971** | **87.766** | **58.511** | **34.479** | **17.383** |

### 4.5 Comparisons on the BindingDB dataset

#### Compared Models

We further assess our model on the BindingDB dataset. We compare our model with four baselines: 1) Similarity-based method Tiresias [Fokoue *et al.*, 2016]; 2) DBN [Wen *et al.*, 2017], a deep learning method using middle-level features from predefined molecular fingerprints and protein descriptors; 3) E2E [Gao *et al.*, 2018] using GCN and LSTM to process drug molecules and protein high-level information (Gene Ontology annotations) respectively; 4) GNN [Tsubaki *et al.*, 2018].

#### Results

The experimental results on BindingDB dataset are demonstrated in Figure 3. Our approach consistently performs well across the test sets and all metrics. Three baselines (Tiresias, DBN, and GNN) perform well on seen proteins, but have much worse performance on unseen proteins. This indicates there exists to be over-fitting over proteins used for training. On the other hand, E2E give a consistent performance for seen and unseen proteins, but it is consistently lower than DrugVQA by 2.6% and 2.3% for AUC and accuracy in average.

**Figure 3:**
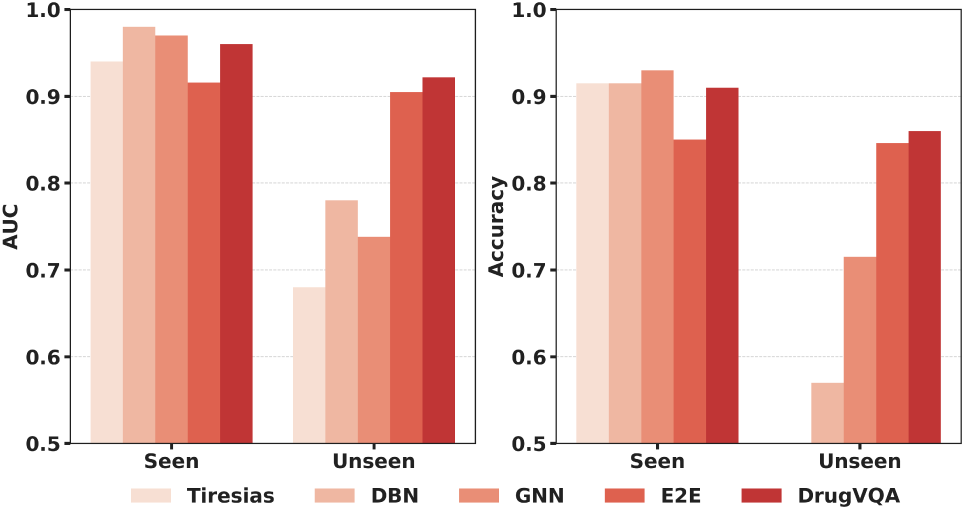
Performance comparisons of our proposed method and baselines on seen and unseen protein targets from the BindingDB.

## 5 Case study

Another advantage of our model is its interpretability. To exemplify this, we selected two top predicted interactions in DUD-E dataset: protein Hsp90 (PDB: 3EKR) and CDK2 (PDB: 2DUV) with their corresponding actives. As shown in Fig 4, the green highlighted the sites with high attentions in the binding pocket, and the red cloud indicates drug atoms with attentions. Darker colors indicate higher attention coefficients. In both cases, molecular components with weights higher than 0.6 overlap substantially with the interaction sites between a molecule and a protein. In the meanwhile, for Hsp90 (left), the pocket importance map highlights residues Asn51A, Asp93A, Met98A, which highly overlap with the key pocket residues observed in the co-crystal complex (PDB: 2DUV). For CDK2 (right), the highlighted key residues (Phe80A, Asp145A) and ligand functional groups in the importance maps show high similarity to observed interactions in the 3EKR. This result suggests that our model can provide reasonable cues for drug-protein binding mode, which is helpful for finding promiscuous domains and designing the active improved drug compounds.

**Figure 4:**
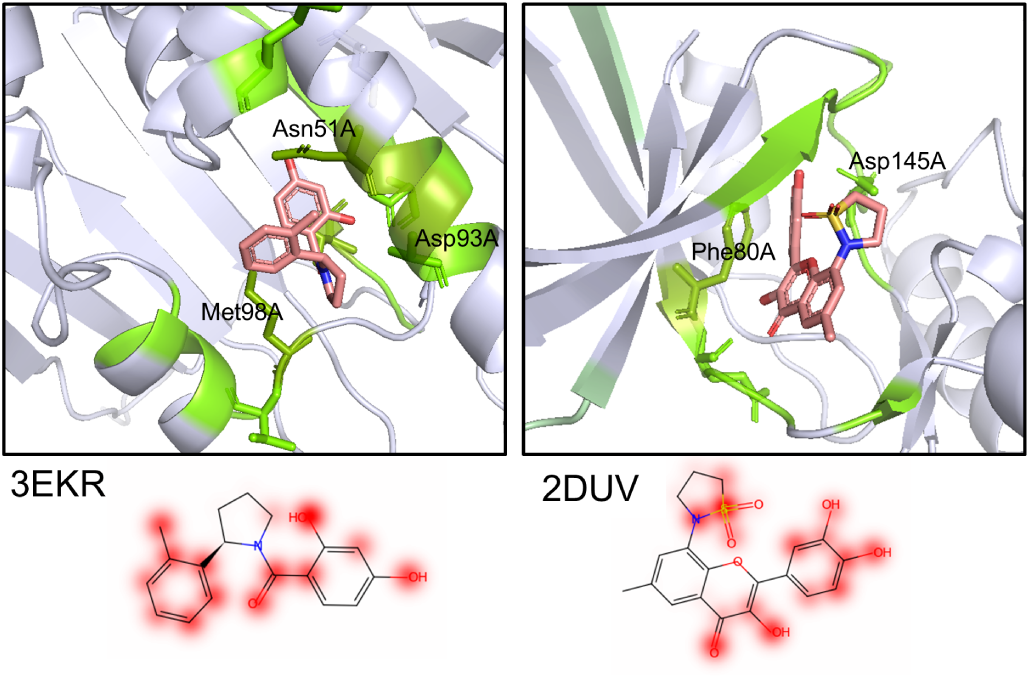
Importance visualization of pocket and ligand pairs (PDB ID:3EKR, 2DUV). The green highlighted the sites with high attentions in the binding pocket, and the red cloud indicates drug atoms with attentions. Darker colors indicate higher attention coefficients.

## 6 Conclusion

In this article, we have presented a novel end-to-end deep learning framework like Visual Question Answering (VQA) task to predict drug-protein interactions. It is the first time to employ self-attentive convolutional and recurrent structures for extracting features simultaneously from protein 2D distance map and molecular language in DPI study. Experimental evaluations demonstrate that our model consistently shows the best performances on three public datasets. Furthermore, the model is shown to be able to provide biological insights for understanding the nature of molecular interactions.

## Acknowledgments

This work has funded in part of the national science & technology major project of the ministry of science and technology of China (2018ZX09735010), and Natural Science Foundation of China (U1611261 and 61772566).

## Footnotes

1 https://deepmind.com/blog/alphafold/

